# Individual identification of Japanese macaques *(Macaca fuscata)* using a face recognition system and a limited number of learning images

**DOI:** 10.1101/2020.07.12.199844

**Authors:** Yosuke Otani, Hitoshi Ogawa

**Affiliations:** CO Design Center, Osaka University, Osaka, Japan; Faculty of Information Science and Engineering, Ritsumeikan University, Shiga, Japan

**Keywords:** Deep learning, face and facial feature detection, face identification, machine learning, wildlife monitoring

## Abstract

Individual identification is an important technique in animal research that requires researcher training and specialized skillsets. Face recognition systems using artificial intelligence (AI) deep learning have been put into practical use to identify in humans and animals, but a large number of annotated learning images are required for system construction. In wildlife research cases, it is difficult to prepare a large amount of learning images, which may be why systems using AI have not been widely used in field research. To investigate the development of a system that identifies individuals using a small number of learning images, we constructed a system to identify individual Japanese macaques *(Macaca fuscata yakui)* with a low error rate from an average of 20 images per individual. The characteristics of this system were augmentation of data, simultaneous determination by four individual identification models and identification from a majority of five frames to ensure reliability. This technology has a high degree of utility for various stakeholders and it is expected that it will advance the development of individual identification systems by AI that can be widely used in field research.

## Introduction

Individual identification is a highly important technique in animal research because observing specific individuals across days, months, and years informs use about various events such as relationships among individuals, detailed life histories, individual behavioral differences, and reproductive success (Caro 1998). This technique has been used to study a wide range of mammals, including dolphins, giraffes, elephants, and leopards (Wells and Scott 1990; Wittemyer 2001; Jackson et al. 2006; Muller 2018). However, because individual identification of recorded or remembered characteristics of individuals, such as whisker spots or scars, requires specialized skills (Pennycuick and Rudnai 1970; Friday et al. 2000), there is a limitation that experts must stay in the field for long periods of time to make long-term records. This limitation hinders the progress of individual identified long-term research and the handing over of research. One way to solve these problems is to develop an automated system that can assist in identifying individuals without the need for the specialized skills.

Individual identification systems have been developed, including those with deep learning, that have been used in studying dogs and other domestic animals, penguins, elephants and primates (Burghardt et al. 2004; Loos and Ernst 2013; Kumar and Singh 2014; Allen and Higham 2015; Kumar et al. 2018; Witham 2018; Körschens et al. 2018). However, these studies have been conducted in captive environments where large amounts of learning data are available. At many wildlife research sites, it is difficult to acquire learning images; the number of available learning images per individual is limited or the effort required to acquire images is quite high. Therefore, individual identification methods using AI have not been widely used in field research. By developing systems that can identify individuals from a small number of images, it is possible to expand field research using AI.

The purpose of this study is to investigate the development of a system that can identify animal individuals from a small number of learning images. More specifically, we develop an integrated system that allows us to create a learning model for each group or local population of Japanese macaques. Users can create a training model specific to the target group by loading individually identified images (annotated images) of the target individuals. Although the models obtained by this protocol are not capable of identifying individuals other than the target, the system can be useful for long-term studies of a single or several social groups.

There are two types of errors in individual identification: 1) recognition of a known individual as an unknown individual and 2) mistaking one individual for another. The first error reduces the efficiency of many of investigations that use individual identification, while the second error reduces the accuracy of the investigation. In this study, system development focuses on reducing the ratio of the second error as much as possible and allowing the first error to some extent.

Because our study subject, Japanese macaques *(Macaca fuscata yakui)* in western coastal area of Yakushima Island have been identified and documented by many researchers (Yamagiwa 2008), and multiple neighboring groups were well habituated, the subject were ideal for collecting learning data for individual identification system development and examining the usefulness of that system.

This study is performed following the Research Guidelines for the Study of Wild Primates of the Primate Research Institute, Kyoto University and adhered to the wildlife protection and hunting laws of Japan.

## Materials and methods

### Data collection

Our study subjects were individual Japanese macaques from groups that inhabit the western coastal area of Yakushima. From March 23 - 26 2018, we observed groups that had already been identified and filmed. We obtained 316 videos of 84 individuals for a total of 177 minutes. Of these, 41 videos of 11 individuals from five social groups were used as learning samples (2.45 ± 1.04 min/individual), and 32 videos of the same individuals were used as verification samples (1.22 ± 0.62 min/individual) for individual identification models. Videos taken with different backgrounds were treated as different samples.

### Identification method

There are various methods for human face recognition: principal component analysis using eigenfaces (Navarrete and Ruiz-Del-Solar 2002), linear discriminant analysis (Gorban et al. 2018), elastic bunch graph matching using the Fisherface algorithm (Anggo and La Arapu 2018) and the hidden Markov model (Samaria and Young 1994). Deep learning is effective in identifying many objects because the parameters used for identification are automatically determined. For example, in image processing, systems use image features that cannot be specified in advance by people. Generally, the target of deep learning uses big data and a lot of sample data are required for learning, with more than 1,000 images required for one individual. In this system, in order to obtain a valid identification model from a small number of images, extraction of high-quality learning images and data augmentation were performed. However, since training data are still insufficient, the recognition accuracy of a single model does not achieve sufficient accuracy. We, therefore, integrates four identification models that could each be regarded as a model learned from a different viewpoint by a random function. Each model obtained by performing deep learning several times using the same learning data, which can be regarded as a result from different grounds. Using the Dempster-Shafer theory (Simon 1994; Sentz and Ferson 2002), the results from different grounds can be linked and integrated probabilities can be calculated.

### System Configuration

The system consists of four main components; “data augmentation system” for increasing the number of learning images without quality deterioration, “acquisition system of face detection cascade” for extracting face from images, “learning system of individual identification model” for creating identification models from learning images in which individuals are identified, and “Individual identification system” for determining the individual from the verification image (Fig. 1).

**Fig.1 :**
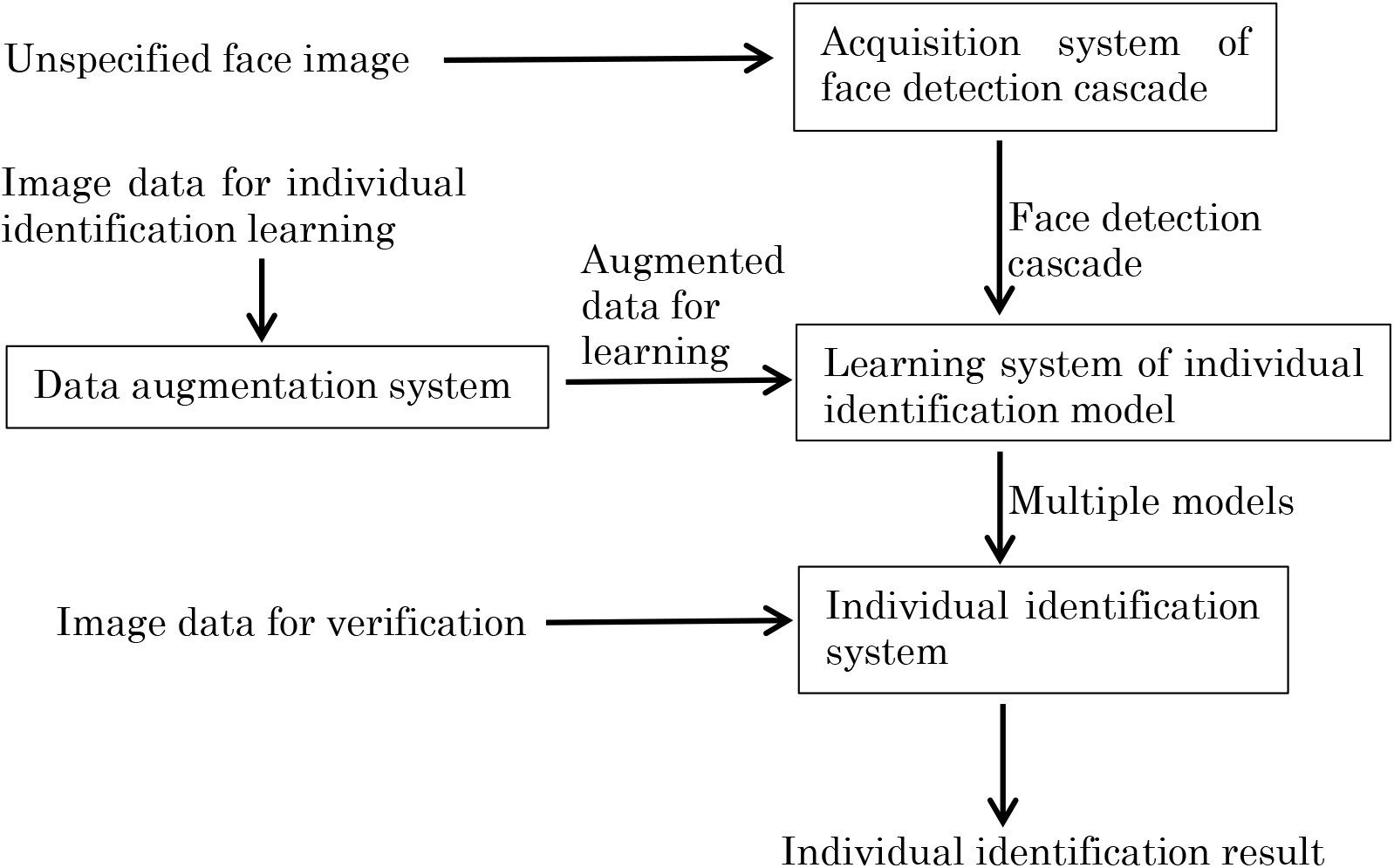
Conceptual diagram of system configuration. Each box represents four main components.

#### Data Augmentation System

This component is for increasing the number of learning images for face detection cascade and individual identification model. Although there are many methods for data augmentation, the following processes that can reduce image quality were not applied: adding noise, brightness adjustment, contrast adjustment, and smoothing. In this study, the following transformations are designed and used for data augmentation:

1. The center of the face area is rotated by 0.1, 0.2 and 0.3 radians to the left and right. Up to six face images are acquired from an image.
2. Compression ratios of 0.95, 0.9 and 0.85 are applied to the vertical and horizontal directions, respectively. Up to six face images are acquired from an image.
3. Compression is also performed after rotation. The rotation and compression rates are the same in (1) and (2), respectively. Up to 36 face images are acquired from an image.
4. Rotation is performed also after compression at the rotation and compression rates that are the same as those in (1) and (2), respectively. Up to 36 face images are acquired from an image.

Examples of the augmented face images are shown in Fig. 2.

**Fig.2 :**
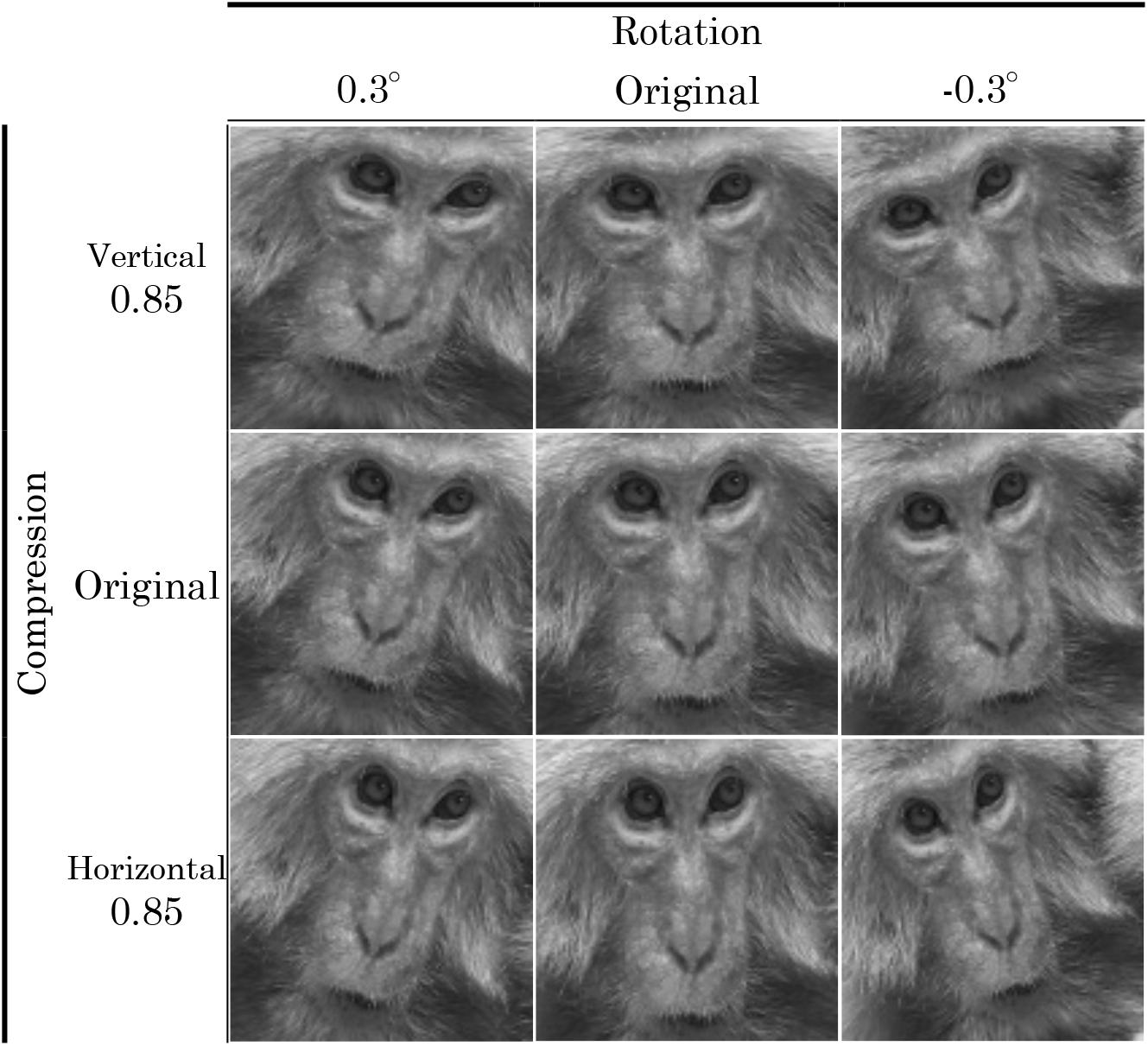
An image obtained by rotating the original image by 0.3 ° to the left and right, respectively, and an image obtained by compressing the original image by 0.85 times in the horizontal direction and the vertical direction are exemplified..

#### Acquisition System of Face Detection Cascade

This component was made to automatically extract face of Japanese macaque from the acquired images (Fig. 3). The OpenCV system provides cascade classifiers (i.e. a classifier-summarizing feature of a series of learning images) based on Haar features (Viola and Jones 2001) such as human faces and eyes, or cat faces. However, the cascade classifier for wild animals including Japanese macaques was not provided and therefore, we built cascade classifiers for Japanese macaque face detection from scratch using the following programs:

1. opencv_createsamples application: creates a file for learning that provides a large set of images with positive samples and location information of the samples.
2. opencv_traincascade application: creates the boosted cascade of weak classifiers based on the positive and negative datasets.

**Fig.3 :**
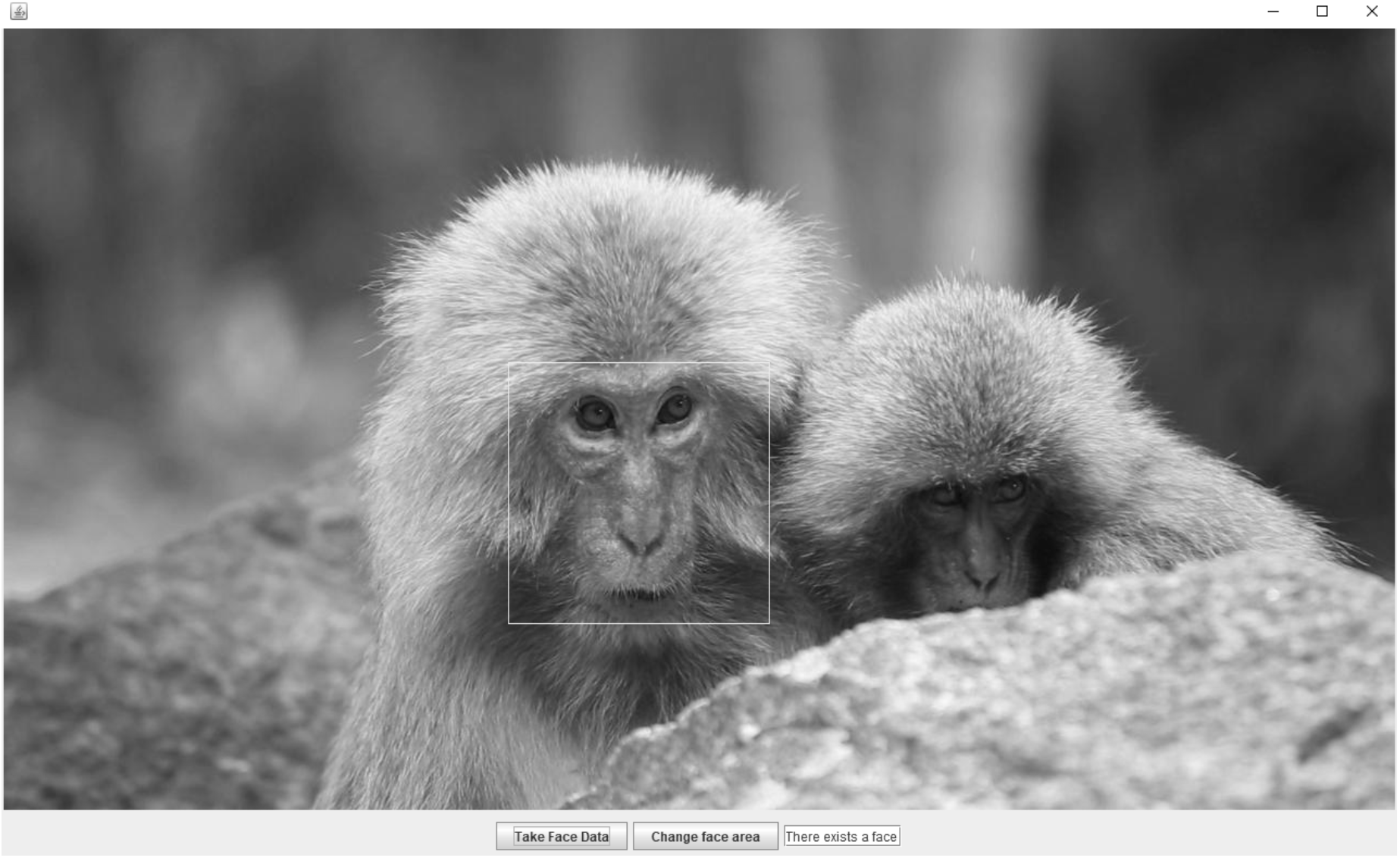
The face of macaques included in the read video is automatically extracted. The white frame indicates the face to be extracted.

To build a face detection cascade, we first manually extracted 224 face images from videos. Using data augmentation methodology, up to 84 new images were acquired from one image. Finally, 23,520 positive images were obtained. As negative controls, 14,790 images other than Japanese macaque were randomly extracted from Caltech 101, which is a publicly available database of digital images (e.g. Fei–Fei and Fergus 2004).

Using the “opencv_createsamples.exe” program provided, a special file (for example, “face.vec”) was created to select positive images at random. Then, a face detection cascade was obtained by the “opencv_traincascade.exe” program. The name of the file (i.e. “face.vec”) of positive images, the number of the positive images (22,000), the number of negative images (14,790), the size of the image area for the cascade (40 by 40), and learning steps (22) served as parameters for the program.

#### Learning System of Individual Identification Model

In this study, TensorFlow (Abadi et al. 2016) is adopted among many deep learning libraries. TensorFlow was developed by researchers and engineers from the Google Brain team (Google LLC) and is the most commonly used software library in the deep learning field. The technical items (as of version 1.13) for TensorFlow include:

1. TensorFlow is a free and open-source software library for dataflow and differentiable programming across a range of tasks.
2. TensorFlow is available on 64-bit Linux, MacOS, Windows and mobile computing platforms, including Android and iOS.
3. TensorFlow has APIs available in Python, JavaScript, C++ and Java.

Convolutional neural networks (CNN) are suitable for learning specific patterns such as images (Goodfellow et al. 2016). In the experiments, the CNN with the following configuration was adopted.

1. Input layer: The color images of the face area are given by 96 pixels in width and 96 pixels in width and 3 channels of data.
2. Convolutional layer: The first, second, third, fourth layer has sixteen 5 × 5 filters, thirty-two 3 × 3 filters, sixty-four 3 × 3 filters and one hundred-twenty-eight 3 × 3 filters, respectively. ReLU (Rectified Linear Units) is used after applying every convolution.
3. Pooling layer: The max-pooling filter that is a square matrix with dimensions 2 × 2 is applied after each convolutional layer.
4. Fully connected layer: The 1023 nodes are used.
5. Output layer: Twelve results are represented using the Softmax function.

The learning rate was 1e-4 and the batch size was 128.

#### Individual Identification System

To improve identification accuracy, four identification models were integrated by Dempster-Shafer Theory (DST) (Simon 1994; Sentz and Ferson 2002) which is a mathematical theory for evaluating evidence. Because the deep learning process involves randomness, even if the learning images are the same, the learning results, that is, the performance and characteristics of the individual identification model, can differ. Therefore, we obtained four individual identification models by learning four times from the same learning images. In DST, evidence can be associated with multiple possible events, such as sets of individuals. A more conclusive finding can be obtained by summarizing the uncertain evidence collected individually. This method has been used to assess the structural safety of existing building and the results were evaluated (Ogawa et al. 1985; Ogawa and Tamura 1986). Since the identify model obtained by learning with this system is not completely reliable for individual recognition, more reliable conclusions can be made using the four identification models.

## Results

### Acquisition of Learning Data for individual identification

Our identification target was 11 individuals form five groups from the western coastal area of Yakushima island. For the program, 12 labels were used for 11 target individuals and non–target individuals which were labeled as “unknown”. To obtain the learning image, an image including a face was used as input, the face area was extracted automatically by *“face detection cascade”* (Fig. 3). When multiple individuals appeared in one image, multiple faces were simultaneously detected and the face of the object individual were arbitrarily selected. From extracted images, those that captured as many facial directions as possible were manually selected as learning images. With this procedure, we acquired 220 images of faces from 41 videos as learning data for constructing the learning system of individual identification model, and increased the image number to 13574 with the data augmentation system (Table I).

**Table I.** Numbers of samples used for learning.

|  | Individual ID |  |  |  |  |  |  |  |  |  |  |
| --- | --- | --- | --- | --- | --- | --- | --- | --- | --- | --- | --- |
|  | ID01 | ID02 | ID03 | ID04 | ID05 | ID06 | ID07 | ID08 | ID09 | ID10 | ID11 |
| **Videos for learning** |  |  |  |  |  |  |  |  |  |  |  |
| Total number of frames | 7522 | 4410 | 6524 | 3752 | 2151 | 443 | 5228 | 5341 | 4559 | 3383 | 5245 |
| Total length (seconds) | 251 | 147 | 218 | 125 | 72 | 15 | 174 | 178 | 152 | 113 | 175 |
| Number of video files | 8 | 3 | 5 | 3 | 2 | 1 | 6 | 3 | 3 | 4 | 3 |
| **Extracted images for learning** |  |  |  |  |  |  |  |  |  |  |  |
| Number of images | 35 | 13 | 33 | 11 | 14 | 5 | 26 | 20 | 12 | 33 | 18 |
| Number of augmented learning images | 2070 | 784 | 1983 | 599 | 858 | 384 | 1474 | 1898 | 832 | 2124 | 568 |

### Performance of the individual identification model

To examine the effectiveness of the system, we loaded videos for verification into the system and calculated the error rate of individual identification. Still images were automatically acquired from the videos and face regions were automatically obtained from still images. For each frame of the video (approximately 30 frames per second), four individual identification models judged the individual and the identification results were determined by a majority decision for five frames of the video. Table II shows the identification results of 11 individuals. In practice, there was an average 19.9% (max 53.8%, min 3.0%) chance that the subject would be identified as unknown, while there was an average 2.2% (max 5.0%, min 0.2%) chance that the subject would be mistaken for another individual.

**Table II.** Results of identification

|  | Individual ID |  |  |  |  |  |  |  |  |  |  |
| --- | --- | --- | --- | --- | --- | --- | --- | --- | --- | --- | --- |
|  | ID01 | ID02 | ID03 | ID04 | ID05 | ID06 | ID07 | ID08 | ID09 | ID10 | ID11 |
| **Videos for verification** |  |  |  |  |  |  |  |  |  |  |  |
| Total number of frames | 2559 | 2079 | 1760 | 3819 | 2085 | 601 | 1269 | 1556 | 3982 | 921 | 3564 |
| Total length (seconds) | 85 | 69 | 59 | 127 | 70 | 20 | 42 | 52 | 133 | 31 | 119 |
| Number of video files | 2 | 3 | 3 | 3 | 2 | 1 | 3 | 3 | 5 | 3 | 3 |
| **Number of samples and correct identifications** |  |  |  |  |  |  |  |  |  |  |  |
| Number of obtained face image sets | 800 | 1408 | 875 | 658 | 1120 | 590 | 338 | 266 | 1578 | 212 | 1765 |
| Number of correct identifications | 648 | 781 | 819 | 636 | 1064 | 457 | 226 | 183 | 1300 | 97 | 1658 |
| **1) error: recognize a known individual as an unknown individual** |  |  |  |  |  |  |  |  |  |  |  |
| Number identified as Unknown | 124 | 556 | 42 | 20 | 54 | 108 | 102 | 76 | 235 | 114 | 94 |
| Rate identified as Unknown | 15.5% | 39.5% | 4.8% | 3.0% | 4.8% | 18.3% | 30.2% | 28.6% | 14.9% | 53.8% | 5.3% |
| **2) error: mistake the individual for another individual** |  |  |  |  |  |  |  |  |  |  |  |
| Number of misrecognitions | 28 | 71 | 14 | 2 | 2 | 25 | 10 | 7 | 43 | 1 | 13 |
| Rate of misrecognition | 3.5% | 5.0% | 1.6% | 0.3% | 0.2% | 4.2% | 3.0% | 2.6% | 2.7% | 0.5% | 0.7% |

## Discussion

In this study, we constructed a system that enables registration and identification of individuals from a small number of learning images. The rate of 2) error which affects the accuracy of the survey was 2.2%. Currently, the number of distinguishable individuals is only 11, but there is a possibility that future development will increase this number with high accuracy by constructing deeper models. Being able to identify a sufficient number of individuals from a small number of images will open the possibility of using AI for individual identification that could be adopted as a general method in many wild animal research sites where it is difficult to obtain a large number of learning images. The characteristics of this system are augmentation of data by rotation and compression, simultaneous determination by four individual identification models, and identification by a majority of five frames to ensure reliability. Additionally, this technique could be applied to the study of other animals.

The current system with a limited number of distinguishable individuals is still useful in active field sites. Even if the number of distinguishable individuals is about 10, it is possible to distinguish social groups by registering individuals that belong to different groups. For example, in an environment where multiple groups of Japanese macaques appear in the same area, it is possible to identify multiple groups by identifying several of the females in each group using this system. Group identification is the first step in many studies of animals that form social groups. If the system allows the researchers to identify the subject groups, the identification process will be much easier because the number of individuals that must be identified is quite limited. In addition, if the system allows non-experts to identify groups, the occurrence of groups can be monitored even when experts are not present.

It is important to note that systems based on a small number of learning images can produce different results at each time of learning. In this study, we have carefully selected the learning images to include faces from many different directions, and if the images were randomly selected, similar results would not be obtained. This paper shows the possibility of achieving a high recognition rate from a small number of images.

Using AI learning systems can enhance manual individual identification in animal populations and will continue to improve over time. For example, the use of video and collected images allows researchers to resume their work after time away from the research site. Moreover, using physical documentation provides continuity as new researchers enter. When a new researcher enters a study site where previous researchers have kept record of individual identification, the researcher must begin by trying to interpreting preserved identification charts of the target group; using a constructed database with AI learning system makes it possible to start and train new researchers efficiently.

The identification system can be used in areas where animal damage occurred. For example, in the case of exterminating Japanese macaques to reduce crop raiding, random elimination may cause division of a group (Uno and Kinota 2019); therefore, selective extermination that specifically identifies individuals that are most harmful to the crop fields is required (Seino et al. 2018). However, selective extermination requires skilled experts and is labor and time intensive. If non-experts can get images of harmful individuals with automatic trap cameras and identify them using AI learning system, it would be possible for residents and local governments to conduct selective extermination. Additionally, the identification system can be used to monitor crop raiding groups. Currently, monitoring is conducted using collar-type telemetry (Yamada and Muroyama 2010; Yamabata et al. 2018) and an approach warning system (Yoshida et al. 2015). However, telemetry is laborious and expensive to install on animals. In conjunction with telemetry, our system could improve crop protection. By combining telemetry that identifies animal location in remote areas and identification system that increases the number of recognizable individuals, effective measures against animal damage to crops or properties can be provided.

It is also considered useful in sightseeing spots. For example, in the wild monkey park, identifying providing information of individuals enables tourists to observe specific monkeys. By providing name, sex, age, mother, personality character, estrus status information etc., it is possible to create new tourist experiences, improve educational effects, and secure macaque and tourist safety.

In the primate research field, many achievements have been made through individual identification, but there are various restrictions associated with this method. By describing individual relationships in detail, the concept of ‘society’ in Japanese macaques has arisen and details have been clarified (Nakagawa et al. 2010). However, such methods can only be applied to animals that can be continuously traced and observed. Deep-learning systems may allow for the identification of more animal species, which would expand the possibilities of wildlife research.

Individual identification techniques in primate research are limited by their dependence on the individual capabilities of the researcher. One of those limitation is number of individuals that a researcher can remember. There is also a temporal restriction. If the investigation is interrupted for several years, it is quite difficult for researchers to keep accurate records during the research hiatus. There is also a problem of data transmission between personnel. In areas where surveys are continuously conducted such as Yakushima, researchers create and preserve identification charts of the survey target groups, which can be used by other researchers. However, although these charts schematically diagram and describe facial features such as the presence or absence of scratches, a training periods is necessary for new researchers to learn how to identify study group individuals. Our identification system was developed with the purpose of reducing the spatial, temporal and capacity limits of conventional individual identification methods. However, with regard to the record of detailed behavior, there is a superiority to the conventional individual identification method and by combining the identification system with the conventional method, it is possible to apply individual identification spreads to the whole research program. Additionally, as a new challenge in primate research, a system has been developed to automatically detect and track Japanese macaque bodies from videos (Ueno et al. 2019). The combination of this tracking system, an identification system and robotics could one day allow researchers to monitor individuals in their lab.

This study enabled the identification of individual Japanese macaques using limited learning images. This technology has a high degree of utility for various stakeholders and will help research advance and develop the field. Further studies are needed to increase the number of identifiable individuals and develop its implementation in various wildlife research fields.

## Acknowledgments

We thank our friends and colleagues in Yakushima for their hospitality and support during the field research, as well as the Yakushima Forest Office and Kirishima-Yaku National Park for granting permission for our study. The Sarugoya-Committee and the Field Research Center of the Wildlife Research Center, Kyoto University provided excellent facilities. We thank the researchers conducting studies in the western coastal area of Yakushima who provided us with information for the identification of Japanese macaques. During the early stage of system development, images of Japanese macaques bred by the Center for Human Evolution Modeling Research in Primate Research Institute, Kyoto university were used, and we want to thank A. Sawada, N Suzuki-Hashido, M. Morimoto, T Natsume, A Kaneko and the members of the center for their support. We thank Drs. G. Hanya and H. Sugiura for their valuable comments. This study was financed in part by JSPS KAKENHI Grant Number 19K15935 to YO, as well as the Cooperative Research Program of Primate Research Institute, Kyoto University.

